# Distribution and diversity of fish species exposed to artisanal fishery along the Sudanese Red Sea coast

**DOI:** 10.1101/763961

**Authors:** Erik Olsen, Bjørn Erik Axelsen, Even Moland, Anne Christine Utne-Palm, Elamin Mohammed Elamin, Motassim Ali Mukhtar, Adel Mohamed Saleh, Sheikheldin Mohamed Elamin, Mohamed Abdelhameed Iragi, Said Gumaa Fadul Gumaa

## Abstract

The semi-enclosed Red Sea harbours one of the longest coral-reef ecosystems on the planet. The ≈ 850 km section of the western shore, comprising the coastline of the Red Sea State of the Republic of Sudan, has however been sparsely studied. Sudan’s coral reef fishery provides livelihoods to fishers and business opportunities by means of local and regional trade, however, the knowledge level of the state of the natural resources and the impacts of fisheries are poorly known. Here we report the results from the first three comprehensive fisheries research surveys spanning the entire Sudanese coast in 2012-13, representing a new baseline for the western coast fisheries resources. The surveys covered the entire coast from inshore down to about 150 m bottom depth using a combination of baited traps, gillnets and handlines to sample the various reef habitats and fish assemblages. The results demonstrate a uniform latitudinal species distribution with peak catch per unit effort rates in and around the Dungonab Bay area in the north and the outer Suakin archipelago in the south. Functional diversity (Rao’s Q index) was found to be highest in and around the Dungonab Bay area, thus coming through as a regional hot-spot of biodiversity. The results form a baseline for future research and monitoring, thus representing key input for an ecosystem approach to management of Sudan’s coastal artisanal fisheries.

## 1 Introduction

With its semi-enclosed location between the African continent and the Arabian peninsula the waters of the Red Sea are warmer and more saline than many other marine tropical ecosystems [1]. The Red Sea is host to a uniquely rich marine biodiversity and high prevalence of endemic species [2,3,4]. While the northern reef areas of Egypt and the Gulf of Aqaba/ Eilat have been extensively investigated [1, 5], the Red Sea proper is generally poorly studied, and only rudimentary studies from decades back have focused on commercial fisheries [6]. The research published on the Red Sea ecosystem is dwarfed by that of the Great Barrier Reef and, in particular, the Caribbean [1], despite equal scientific relevance. The sparseness of information about the fisheries and the state of the resources harvested severely limits the authorities’ ability to sustainably manage the sector in an effective and directed manner [1]. Institutional capacities in the region are limited and official landings data are sparse and essentially unreliable. Tesfamichael and Pauly [6] reconstructed the catch statistics from all Red Sea countries using a combination of unpublished data and interviews with managers, stakeholders and fishers around the Red Sea. They found that the reconstructed catches were 1.5 × larger than the official FAO statistics with artisanal fisheries dominating, accounting for 49% of the total catch from 1950 – 2010 [6]. These estimates, however, also rely on several assumptions and are too rather uncertain, and management and enforcement of the fisheries in the region are essentially based on a highly data-poor knowledge base (Sudan, Eritrea, Yemen) [6].

The Republic of the Sudan’s Red Sea State include 853 km of the 2250 km African (western) Red Sea shore (Figure 1). Although there are large latitudinal gradients in environmental conditions with salinity increasing to the north and temperature increasing towards the south [6, 7], the biological community changes little from north to south [8]. Most studies from the region are over 50 years old and have primarily focused on single reef study sites such as the Costeau Conshelf habitation experiment at Sha’ab Rumi in 1963 [9]. More recently, the Dungonab Bay area north of Port Sudan was studied and deemed to be of global significance and has subsequently been included in the UNESCO list of world heritage sites since 2016 along with the Sanganeb atoll as the “Dungonab Bay – Mukkawar Island Marine National Park” and the “Sanganeb Marine National Park” [4]. Historically, specimen collectors and early natural scientists have described the marine fauna [1,10,11], but thus far there have been no large-scale studies systematically covering the coastal fish assemblages (but see Kattan et al. [7]).

**Figure 1.**
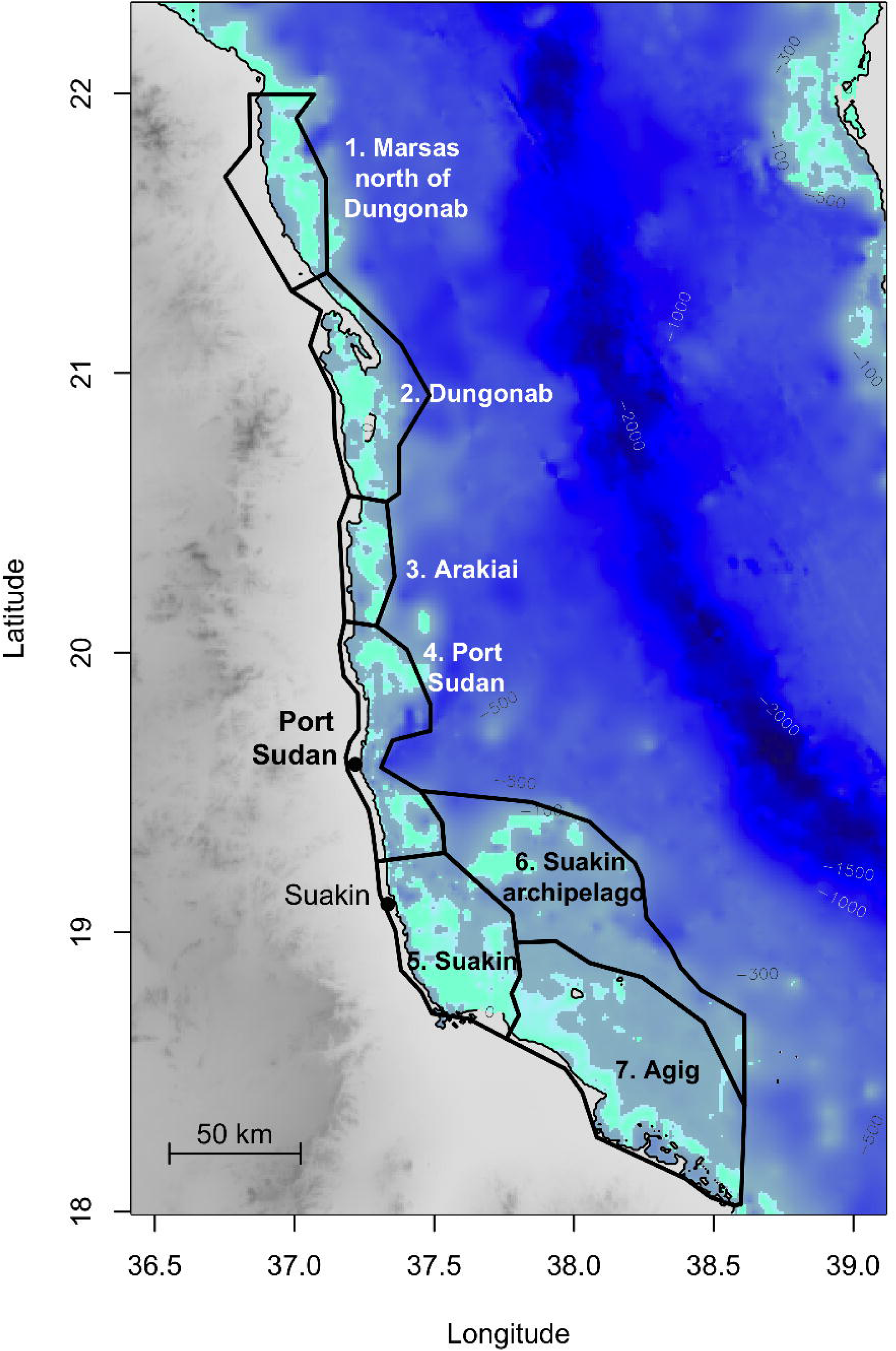
The Republic of the Sudan Red Sea coast, with coral reef areas (aquamarine) and the names and spatial extent of seven fisheries management areas (black polygons) shown.

The 2008 Equipe Costeau expedition was the most comprehensive study to date, but has limited value in terms of describing the marine fishery resources as it only covered a small part of the coast, focusing on local biodiversity rather than abundance [4, 12]. Their teams applied underwater visual census (UVC) methodology to survey the reef fish assemblages and collect benthic data [12]. Kattan et al. [8] carried out comparative UVC surveys of Sudanese and Saudi reefs and found the Sudanese reefs, especially those furthest offshore and to the south, to be in pristine conditions, with biomass levels approaching those observed at remote Pacific islands and atolls. However, the Kattan et al. [8] study did not cover the northernmost section of the Sudanese coast, nor the near shore coastal reef areas.

Several development aid organizations as well as the Red Sea State government have pointed to development in the marine fisheries sector as one possible route to increased revenue, food production and employment in coastal communities. However, an appropriate understanding of the fisheries and its resource base, that can provide guidelines for sustainable harvesting practices and restrictions, is essential in order for such development to be ecologically and economically sustainable in the long term [13].

The data-poor situation as that of the Red Sea State coral reef fishery offers limited opportunity to draw inference regarding the status and evolution of the fishery, the resource base and the sustainability [14]. Despite management efforts in coral reef fisheries worldwide, 55 % of island-based coral reef fish communities are fished in an unsustainable manner [15], and a review of artisanal coral reef fishery research found that nearly 90 % of the studies listed overfishing as a concern [16]. Several strategies have the potential to improve management outcomes, such as strengthening governance, developing a more nuanced understanding of the interaction between fishing, alternative livelihoods and human well-being, and explicitly linking gear selectivity to ecosystem effects [17]. A recent study has raised concern that the roving coral grouper *Plectropomus pessuliferus marisrubri* and the squaretail coral grouper *P. areolatus* (Arabic names ‘Najil’ and ‘Silimani’, respectively), both highly valued target species in the hook-and-line fisheries, are already in an overfished state [18], a situation documented on the eastern, Saudi Arabian, side of the Red Sea [8, 19].

### BOX – Sudan’s coral reef fisheries

The marine fisheries sector in Sudan is comparatively small with annual catches reported at 5700 tons according to the official FAO statistics [21, 22], while according to the catch reconstruction of Sudanese fisheries [23] the annual catch for 2010 was 2000 tonnes, predominantly from artisanal handline- and gillnet fisheries on and along the fringes of coral reefs operating from many small landing sites and villages along the coast. The management of the artisanal fishery is divided into seven management areas (Figure 1). Tesfamichael and Elawad [23] attributed the discrepancies between the official catch statistics and the reconstructed catches to misreporting on the side of the Sudanese fisheries authorities.

The artisanal fishing fleet currently consists of approximately 2000 fishers operating about 1000 vessels that are mostly in the range of 6-10 m of length, many equipped with 30–40 horsepower outboard engines (Marine Fisheries Administration, unpublished data). Most of these vessels target fin-fish (particularly the high-priced groupers) and have facilities for storage of ice and catch. With a crew of typically 2-5 fishers these vessels operate throughout the near- and offshore reef systems and archipelagos for several days up to two weeks per fishing trip. The most common fishing gear is handline, but mono- and multifilament gillnets are also prevalent in some areas. Some of the larger vessels bring small flat-bottom dories to reach into lagoons and onto reef flats to deploy gillnets (barrier nets) for capturing roving herbivores (parrotfish, surgeonfish) that are chased into the nets by fishers using snorkelling gear.

Sudan is the Red Sea country that utlizes its marine resources to the least degree [8] and compared to other countries bordering the main part of the Red Sea, Sudan has the lowest catch per km of coastline [6], governing some speculation that there may be room for expanding fisheries yields.

### 1.1 Aims

The project “Surveys of renewable marine resources in the Red Sea State” was implemented in the period from 2012 to 2013 [24] with the main aim to provide a baseline study of the available fish resources along the entire coast. Three monitoring surveys using different fishing methods were carried out, at two at different periods of the year, and in two subsequent years. Because institutional and technical capacity in Sudan was limited the projected opted to employ survey and monitoring methods that required as little technology and resources as possible while still gathering quality data. This approach thus excluded diver-based methods like UVC that requires extensive training, expensive equipment and costly maintenance of equipment.

During each survey, conventional fishing methods such as gillnets and handlines were operated in conjunction with baited traps to sample the proportion of fish assemblages vulnerable to these fishing gear types, thus enabling comparison of the efficiencies of various fishing methods for monitoring fish assemblages in coral reef areas. The project also provided training opportunities for local scientists, managers and students in fisheries research and monitoring methods.

Here, we report on the catch rates, species densities and functional diversity from trap, gillnet and handline catches obtained during three surveys conducted in November 2012, May 2013 and November 2013 covering all seven management regions of the Sudanese coast (Figure 1). Here we document a baseline of the fisheries resources along the coast of Sudan and provide updated information on biodiversity hot spots, distribution and indices of relative abundance. The results provide valuable basic biological information on several Red Sea fish species that are currently unavailable in the public domain (FishBase) [24], and are relevant with regards to the ecological understanding of fish communities in the Red Sea

## 2 Material and Methods

The sampling scheme was stratified according to the seven established fisheries management areas (Figure 1). The Sudanese coastal shelf is narrow, especially in the north, before it drops off into the deep waters of the central Red Sea. Extensive fringing near-shore coral reefs extend over much of the shelf area complicating both navigation and anchoring, effectively restricting surveying to well-established sea-ways during daytime, consequently producing a somewhat patchy sampling design (Figure 2).

**Figure 2.**
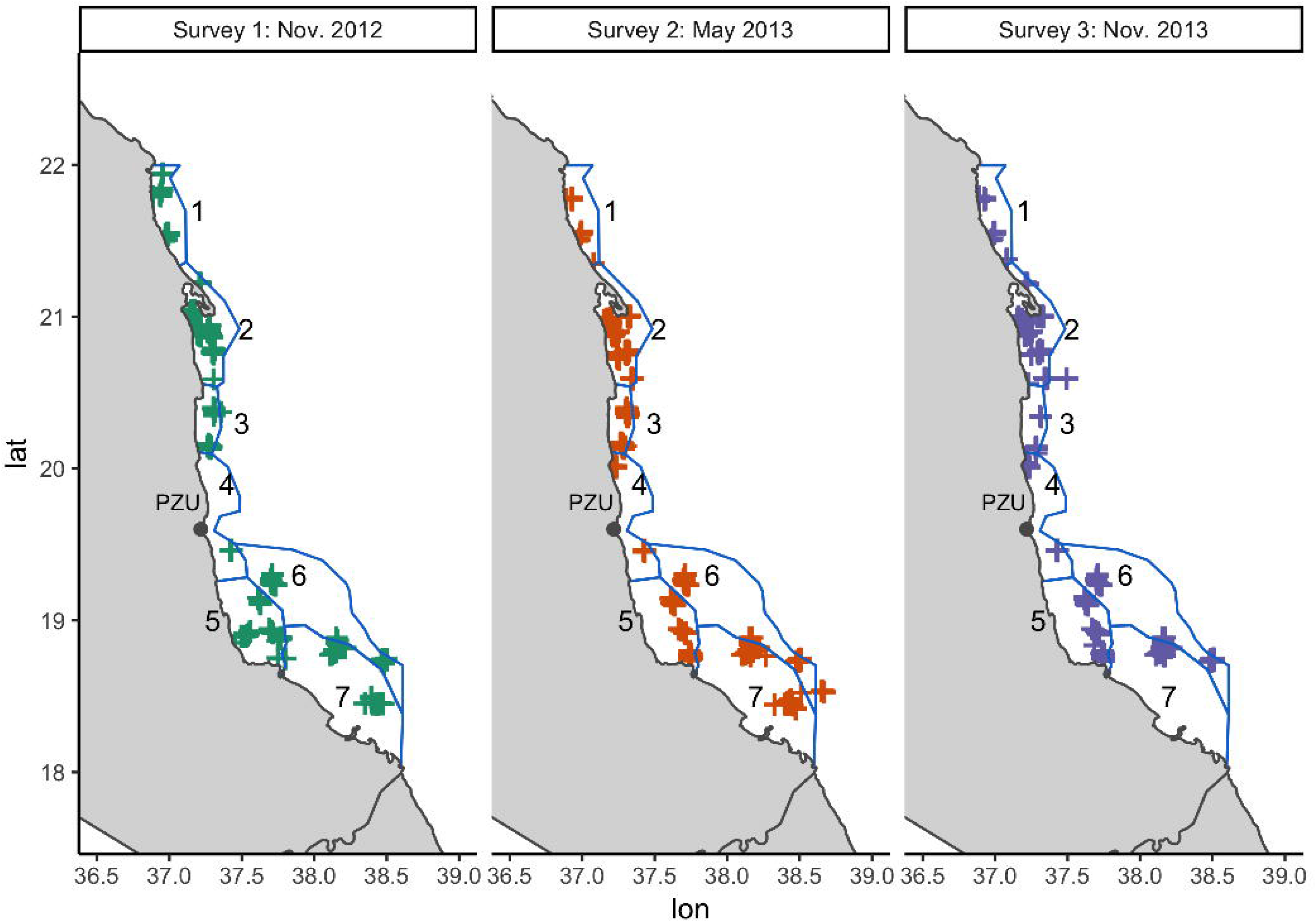
Map showing all survey stations sampled from 2012 – 2013 overlaid the seven fisheries management areas. PZU: the city of Port Sudan.

The surveys covered the coast from Egypt in the north to the border with Eritrea in the south. Two vessels were used: the M/S ‘Don Questo’ (32 m LOA), a previously British oceanographic research vessel currently equipped for liveaboard diving, was used as base for measurement and sampling of the catch, accommodation and meals, and a sheltered fiberglass vessel (10 m LOA) with an inboard engine was used to deploy and retrieve traps and for oceanographic measurements.

The same locations were sampled on each cruise in the northern and central regions to ensure comparability between surveys, producing similar geographical coverages there (Figure 2). In the southernmost part of the study, however, challenging weather conditions restricted the degree of coverage and in November 2013 the survey had to be cut short due to technical delays. The reef at Habily Lory was only sampled in May 2013, while the reef location at Abu Marina was not sampled in November 2013

### 2.1 Fishing gear

In addition to the fishing gear used conventionally by Sudanese fishers (handlines and gillnets), an alternative approach using collapsible baited fish traps was employed to obtain samples from the coral reefs with minimum damage to coral colonies and to target species with low catchability using the artisanal handline method.

#### Baited traps

The main fishing gear used were collapsible fishing traps baited with sardines. The traps measured 150 × 180 × 80 cm and were constructed from steel frames with plastic coated square steel mesh with approximately 50 mm bar length. The number of traps deployed at each reef area ranged from 5 to 14, depending on the site-specific geography (topography and reef length) and weather conditions (wind strength and wave heights) at the time when traps were deployed. The traps were deployed at a mean depth of 21.5 m, but depth varied from 5 – 145 m between areas and surveys (Figure 3). Mean set depth never exceeded 50 m, but deeper traps were set in all areas except area 5 in November 2012, with the deepest traps set at 142 m in area 1 in Nov. 2012 and at 145 m in area 2 in May 2013 (Table 1, Figure 3). Area 5 had the shallowest distribution of trap set depths (and for May 2013 and Nov. 2013 the shallowest mean trap set depth), consistent with this area being inshore of the outermost reefs hence having a shallower average topography than the other management areas. When using traps fish were caught alive and in good conditions, thus any sharks, moray eels or other red-listed species were immediately released upon retrieval of the gear. Whenever caught, species ID and approximate length was noted prior to their release.

**Figure 3.**
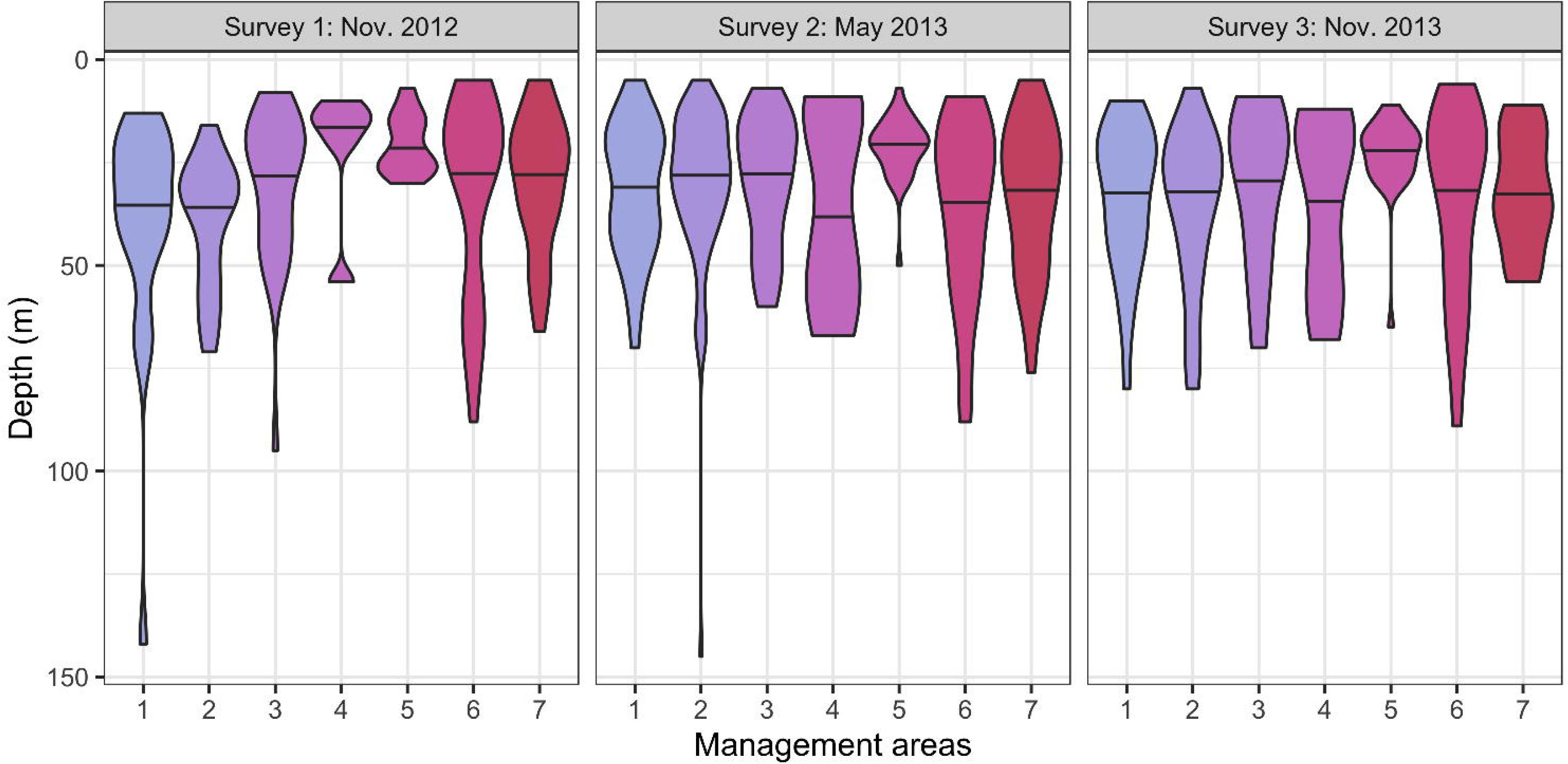
Violin plots of the depths at which traps were set during the three surveys (Nov. 2012, May 2013 and Nov. 2013) in each of the seven management areas (see. Figure 1). The width of the violins are scaled to the relative number of traps set at each depth. Horizontal line in each violin plot marks the mean depth.

**Table 1.** Number of gear deployed (fishing stations) in the seven management areas, total fishing time in each management area for each survey, and minimum, mean and maximum fishing depth for traps for each area and survey. Surveys were carried out in the coastal areas of Sudan in November 2012, May 2013 and November 2013

| Survey | Mgmt. Area | Number of stations (gears set) |  |  | Hours fishing |  |  | Fishing Depth |  |  |
| --- | --- | --- | --- | --- | --- | --- | --- | --- | --- | --- |
|  |  | Traps | Handlines | Gillnets | Traps | Handlines | Gillnets | Average | Maximum | Minimum |
| Nov. 2012 | 1 | 22 | 0 | 0 | 694 | 0 | 0 | 42 | 142 | 13 |
|  | 2 | 54 | 3 | 3 | 722 | 78 | 62 | 41 | 71 | 0 |
|  | 3 | 26 | 0 | 1 | 451 | 0 | 14 | 31 | 95 | 8 |
|  | 4 | 5 | 0 | 0 | 77 | 0 | 0 | 23 | 54 | 10 |
|  | 5 | 31 | 0 | 4 | 678 | 0 | 78 | 21 | 30 | 7 |
|  | 6 | 36 | 0 | 1 | 850 | 0 | 38 | 32 | 88 | 0 |
|  | 7 | 31 | 0 | 8 | 712 | 0 | 136 | 31 | 66 | 5 |
|  | Sum | 205 | 3 | 17 | 4184 | 78 | 328 |  |  |  |
|  | W.catch | 109 | 3 | 12 |  |  |  |  |  |  |
| May 2013 | 1 | 29 | 0 | 1 | 420 | 0 | 24 | 31 | 70 | 5 |
|  | 2 | 81 | 3 | 5 | 1102 | 378 | 222 | 27 | 145 | 5 |
|  | 3 | 32 | 0 | 1 | 546 | 0 | 24 | 29 | 60 | 0 |
|  | 4 | 13 | 0 | 10 | 160 | 0 | 138 | 30 | 67 | 9 |
|  | 5 | 33 | 2 | 5 | 209 | 192 | 196 | 20 | 50 | 7 |
|  | 6 | 45 | 1 | 2 | 642 | 156 | 78 | 35 | 88 | 9 |
|  | 7 | 39 | 2 | 0 | 666 | 186 | 0 | 33 | 76 | 5 |
|  | Sum | 272 | 8 | 24 | 3746 | 912 | 681 |  |  |  |
|  | W.catch | 141 | 8 | 16 |  |  |  |  |  |  |
| Nov. 2013 | 1 | 23 | 1 | 4 | 272 | 3 | 84 | 30 | 80 | 10 |
|  | 2 | 57 | 2 | 6 | 500 | 111 | 156 | 38 | 80 | 7 |
|  | 3 | 9 | 0 | 2 | 123 | 0 | 36 | 27 | 70 | 9 |
|  | 4 | 16 | 2 | 4 | 151 | 5 | 143 | 33 | 68 | 12 |
|  | 5 | 30 | 2 | 3 | 317 | 7 | 147 | 26 | 65 | 11 |
|  | 6 | 40 | 4 | 2 | 444 | 32 | 72 | 40 | 89 | 6 |
|  | 7 | 22 | 0 | 2 | 172 | 0 | 203 | 35 | 54 | 11 |
|  | Sum | 197 | 11 | 23 | 1979 | 157 | 841 |  |  |  |
|  | W.catch | 62 | 9 | 23 |  |  |  |  |  |  |

#### Gillnets

Pelagic gillnets were deployed at dusk and hauled at dawn, recording the exact time of setting and hauling. The nets were inspected hourly to release any sharks or turtles caught inadvertently, after the species, approximate length and catch position had been recorded. Turtles, however, were not identified or recorded in the data. Two types of pelagic gillnets were used at each site: two multi-monofilament gillnets of 28.0 m length and 10.5 m height with 89 mm mesh size (stretched), and one multifilament gillnet of 40.0 m length and 10.5 m height with 76 mm mesh size. The gillnets were anchored to the bottom at one end, with a float attached at both ends, and smaller floats running along the float line to ensure that it floated at the surface. Nets were deployed in channels between reefs, or in open waters inshore or off-shore of the reefs. At some particular shallow stations, the gillnets reached from the surface to the bottom.

#### Handlines

On some locations, during daytime, simple monofilament handlines with a single hook (size 5 – 6) and lead sinker were fitted with fresh bait (sardines) and fished by 2-3 fishermen. The fishermen were told to fish for a standard period of two hours and record the time they started and stopped fishing. However, this information was not well communicated, and for most locations the fishermen returned the next day with the catch but lacking records of how long they had been fishing. Therefor the actual duration of the fishing trials was uncertain and resulting catches could thus only be used for composition of the catch by gear type rather than catch rates as anticipated.

In total 760 stations were sampled during the three surveys (Table 1). The number of gear units deployed increased during the second and third survey because the participants had gained experience and worked more efficiently. Traps were the most common gear with 674 sets. However, 25 % (220 traps) were empty when hauled, compared to only 5 % of gillnet hauls being empty. Use of gillnets increased from 34 in the first survey to 137 in the November 2013 survey, while the use of handlines and traps remained more constant (see Table 1).

### 2.2 Biological measurements

Upon gear retrieval fish caught were removed and placed in numbered plastic boxes on the deck of the vessel where they asphyxiated, as is the normal practice in fisheries operations throughout history as well as in modern fisheries research surveys. This is, however, seldom specified in articles describing fisheries research surveys (see [26–28] for examples).

The length-weight relationship for sharks and moray eels, which were released immediately after capture, was estimated from published length-weight relationships [25]. All other species caught were brought to the ‘Don Questo’ where they were identified to species, and their total lengths and total wet weights were recorded.

### 2.3 Calculation of catch-per-unit-effort

Assuming that the catch coefficient remains constant, the catch-per-unit-effort (CPUE) is proportional to abundance [6], although with caveats regarding hyperstability (CPUE remains stable while abundance is declining) and hyperdepletion (CPUE declines more than the actual decline in abundance) [29]. To facilitate relative comparisons between surveys and gear, CPUE for each species group were standardized as kg fish caught per hour of fishing.

The catch data contained many stations with no catch resulting in CPUE data being non-normal, and rather zero-inflated. Variabilities in CPUE’s between areas, surveys and by depth were therefore evaluated using a zero-adjusted GAM model with depth, area, survey and gear as dependent factors, calculated using the “GAMLSS” package in R. Two models, including or excluding gear as a factor, were evaluated. As the AIC were comparable (−1080 and −1082) we chose to use the model including gear as a factor as this would yield information on the potential effect of gear on CPUE.

### 2.4 Species diversity and functional diversity

Species caught were classified according to the species traits of Stuart-Smith et al [30]: trophic group (carnivores, herbivores, corallivores, invertivores, planktivores), trophic level, maximum length, place in water column, diel activity, habitat and gregariousness. Following the methods of Stuart-Smith et al. [30], we evaluated species density and functional diversity to delineate biodiversity of the catchable component of coastal fishes along the Sudanese coast. Estimation of biodiversity measures were done using the catch data from traps only as neither the gillnets nor handlines covered all management areas during all surveys (see Table 1).

Species densities were calculated as the number of species caught at each station area per hour of fishing, and averaged by management area,. Variability in species density between stations was evaluated using a zero-adjusted GAM model with depth, area and survey as dependent factors, and calculated using the “GAMLSS” package in R.

The Rao’s Q functional group richness index was calculated using the ‘dbFD’ function in the ‘FD’ package in R using all species occurring in three or more traps aggregated across all surveys applying ‘lingoes’ correction to achieve an Euclidian species by species distance matrix. Dimensionality was limited to 10 PCoA axes to avoid integer overload during calculations. The quality of the reduced vector space representation of the traits, *R*^2^, was 0.42 with 10 PCoA axes, using all traits. To evaluate the contribution of individual trait on the functional diversity the model was rerun seven times excluding one trait at a time. As the *R^2^* was lower for all models excluding a trait, except when excluding maximum length, when *R^2^* increased to 0.468, an increase of 0.047, the model excluding maximum length was adopted. This observed increase in *R^2^* when excluding maximum length was likely caused by the large spread in maximum length across the species caught, with few catches of species with a maximum length > 200cm (mostly sharks, morays and large Scombridea).

### 2.5 Data management, statistical and GIS analysis

All data from the surveys were entered into a NAN-SIS [31] database. The tropic group of species in our samples was added to the catch data from a list of traits of coral reef fish from [30], amended with information from FishBase [25] for the Red Sea species not covered in the original species traits list. Detailed information on the position, depth, date, and time, for all fishing stations and all three gear-types as well as species-specific catch information was recorded in the data files (station.csv and catch.csv) deposited on GitHub (https://github.com/erikjsolsen/Sudan). All plotting and statistical analysis was carried out using the R Statistical software package version 3.5.2 (Eggshell Igloo) [32]. The scripts used for all plots and analyses are also made available on GitHub.

## 3 Results

### 3.1 Catch composition by gear

The catch composition of the three fishing gears was markedly different (Figure 4, A), with the gillnets being the only gear catching species in the Chirocentridae family. Trap catches were dominated by Serranidae, Lutjanidae, Lethrinidae and ‘other species’. Gillnets were the most efficient gear towards Carangidae and Scombridae but caught relatively few Lutjanidae and Lethrinidae species that made up the majority of the handline and trap catches. Serranidae were rare in the handline catches, except in May 2013, contrasting the prevalence of larger bodied serranid groupers in the artisanal handline fishery. In terms of trophic groups (Figure 4, B), carnivores dominated the gillnet catches and was the most important trophic group in the trap catches. Invertivores constituted the second largest catch of the traps, while planktivores were the second largest catch of the gillnets. Hand-lines only caught carnivores and invertivores, of which the former dominated the catches.

**Figure 4.**
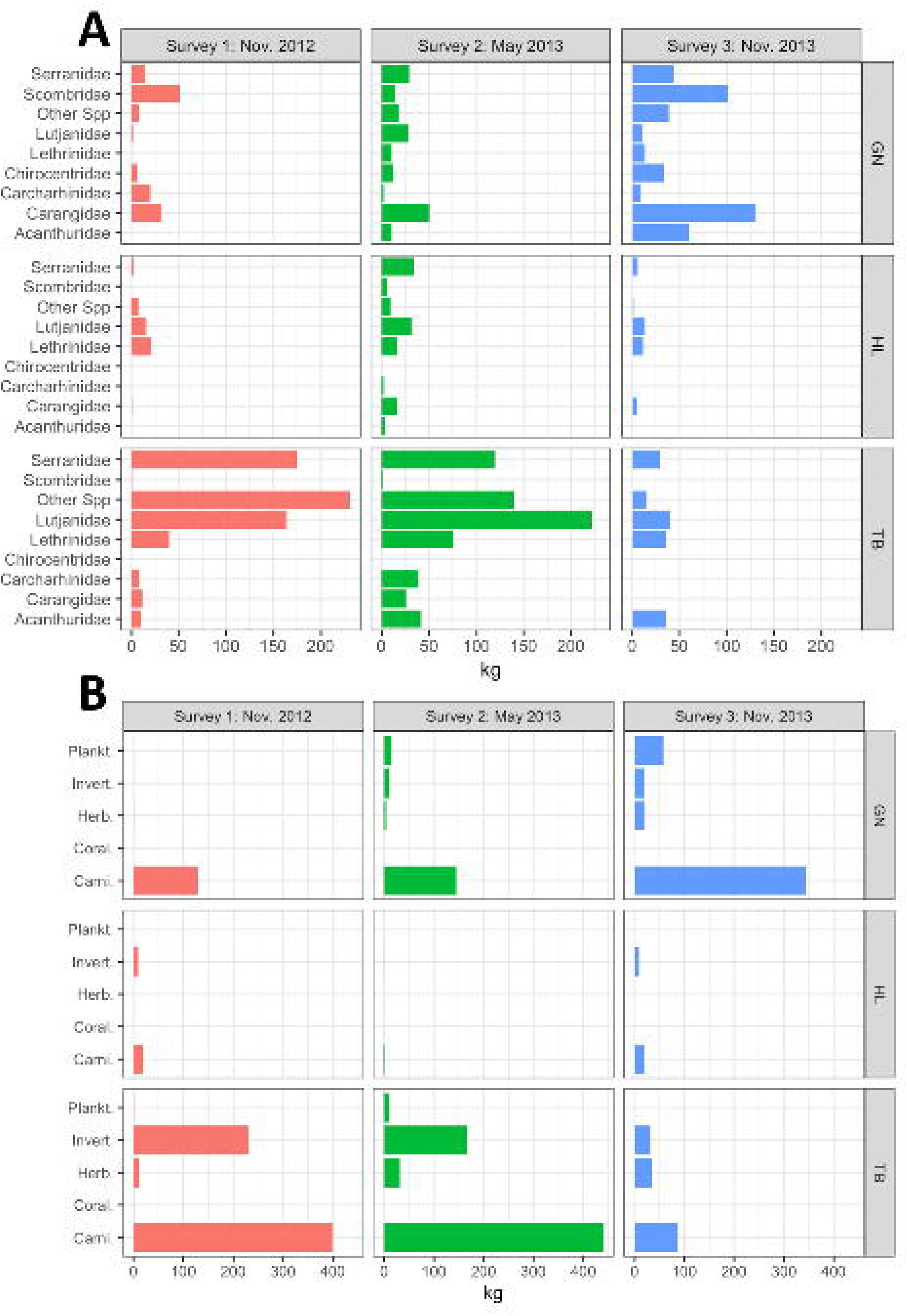
Total catches (kg) for the three fishing gear types: Gillnet (GN), Handline (HL) and Traps (TB), for each of the three surveys (Nov. 2012, May 2013 and Nov. 2013 split according to: (A) main fish families, and (B) trophic group

### 3.2 Cath per unit effort (CPUE)

#### 3.2.1 Effect of gear, survey, area and depth on CPUE

Catch rates between management areas showed large variabilities (Figure 5). Evaluating the combined effects of gears, depth, management area and survey on the biomass CPUE in a zero-adjusted Gamma GAM model (AIC: 307, df:14,– see SI Table 1 for a full summary of model analysis) showed that all gear types, survey (November 2013), and area (areas 1, 5, 6 and 7) had a significant effect, while depth did not have a significant effect on catch rates. Area 2 had the highest recorded CPUE in all three surveys, while area 3 had the highest average CPUE in Nov. 2012, area 6 in May 2013 and area 6 in Nov. 2013 (Table 2). Relative difference in catches of the different family groups varied between the surveys (see Figure 4).

**Figure 5.**
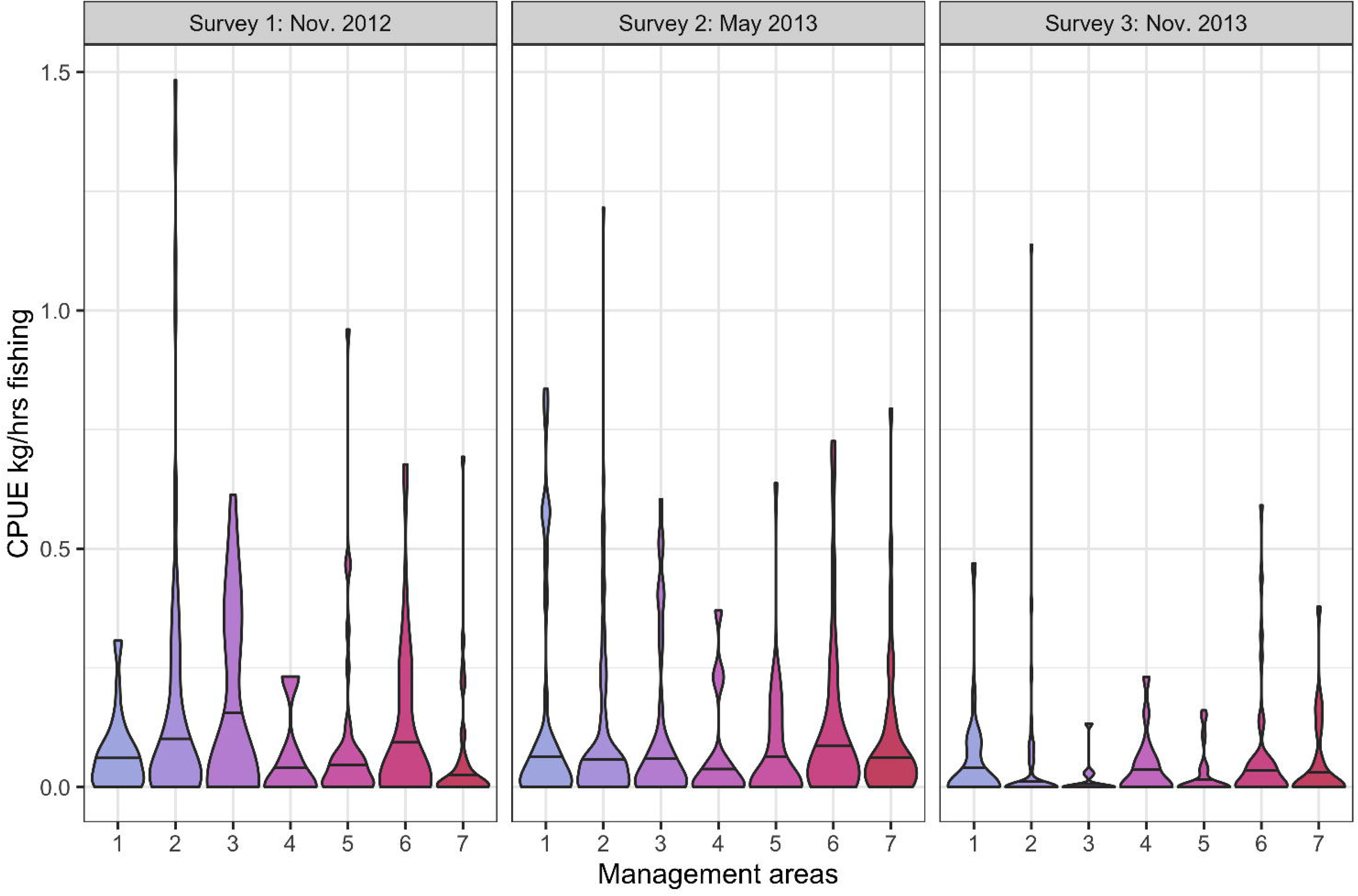
Violin plots of catch rates (kg per hour fishing) for traps in each of the seven management areas and each of the three surveys. The width of the plots are scaled to the relative frequencies of catches with that catch rate.

**Table 2.** Mean CPUE (kg per hours of fishing), mean trophic level and gregariousness of traps and gillnets. Species density (no species in traps per hour of fishing) and the functional group richness of trap catches (Rao’s Q) for each of the seven management areas along the Sudanese Red Sea Coast.

| Mgmt.<br>Area | CPUE (kg/hrs) |  | Trophic level |  | Gregariousness |  | Species<br>Density<br>(traps) | Rao's Q<br>(traps) |
| --- | --- | --- | --- | --- | --- | --- | --- | --- |
|  | Traps | Gillnets | Traps | Gillnets | Traps | Gillnets |  |  |
| 1 | 0,12 | 1,64 | 3,83 | 3,93 | 1,51 | 2,41 | 1,07 | 0,051 |
| 2 | 0,15 | 1,18 | 3,82 | 3,89 | 1,48 | 2,26 | 2,20 | 0,079 |
| 3 | 0,14 | 0,31 | 3,84 | 4,14 | 1,55 | 2,50 | 1,44 | 0,074 |
| 4 | 0,06 | 0,70 | 3,77 | 4,28 | 1,36 | 2,11 | 0,71 | 0,048 |
| 5 | 0,08 | 0,73 | 3,80 | 3,85 | 1,63 | 2,18 | 1,12 | 0,073 |
| 6 | 0,14 | 0,27 | 3,76 | 3,93 | 1,53 | 2,29 | 1,39 | 0,048 |
| 7 | 0,10 | 0,68 | 3,83 | 3,61 | 1,57 | 1,90 | 1,45 | 0,064 |

#### 3.2.2 Trophic group composition of catches for traps

The trap catches (Figure 6) were dominated by the carnivores in all areas during all surveys, except during the Nov. 2012 survey for area 3 where the invertivores had higher CPUE. Corallivores were only caught during the Nov. 2012 survey in areas 2 and 6, while herbivores were caught in all surveys, with the highest catch rates in area 2 and 6. Invertivores had the second highest catch rates, present in all surveys, but with highest CPUE in the Nov. 2012 and May 2013 surveys. Planktivores were caught at very low CPUE in area 7 during the Nov. 2012 survey, but in May 2013 they were also caught (at low CPUE rates) in areas 2, 3, and 5 (but not in area 7), while in Nov. 2013 they had the second highest catch rate in area 2 (after the carnivores) and were also caught in areas 5 and 6.

**Figure 6.**
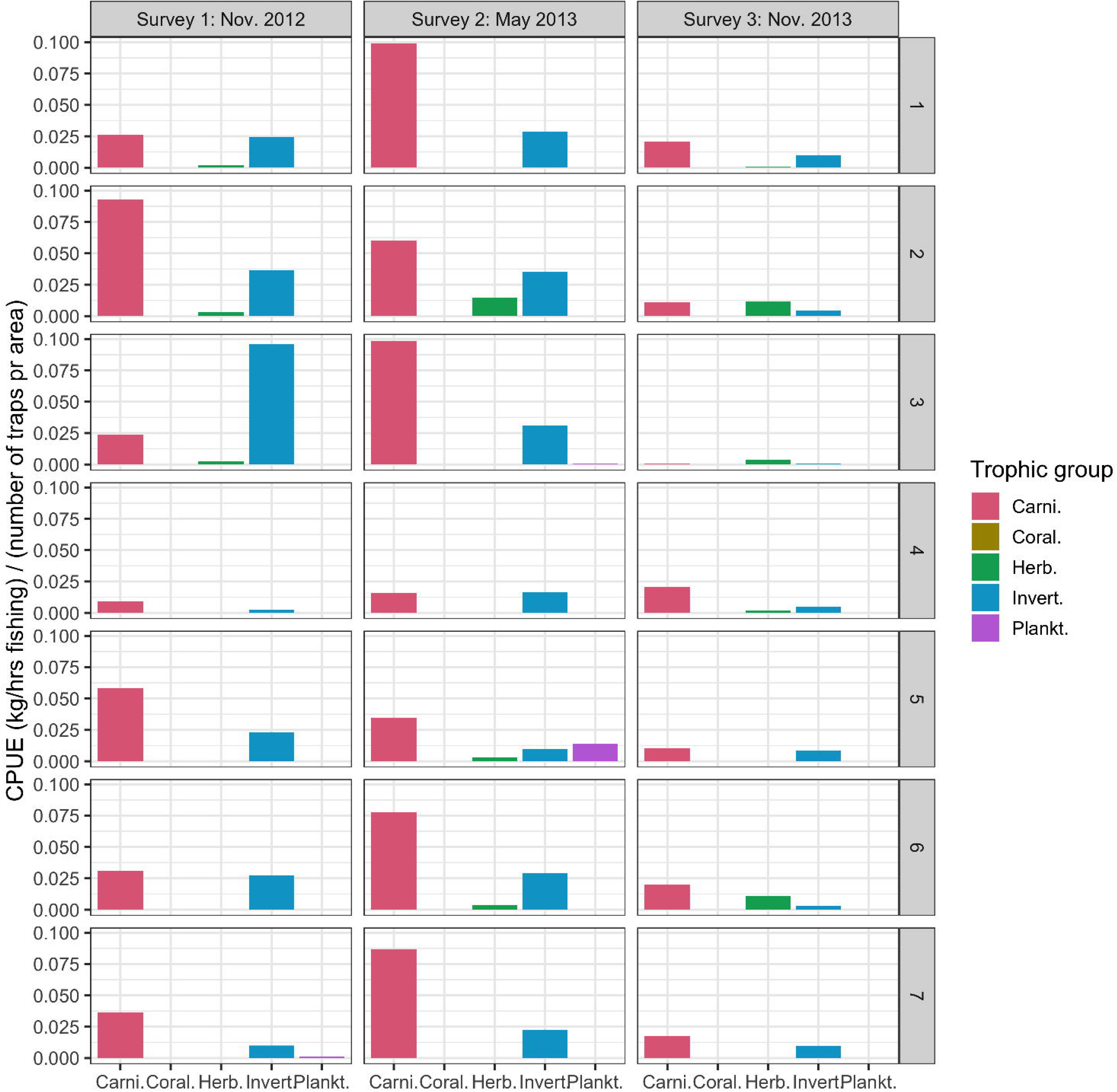
Cumulative catch rates (kg per hour of fishing) of carnivores (Carni), corallivores (Coral.), herbivores (Herb.), invertivores (Invert.) and planktivorous (Plankt.) fish, per trap for each management area and survey by the trophic groups: carnivores, corallivores, herbivores, invertivores and planktivores.

#### 3.2.3 CPUE of trophic groups by depth for traps

Carnivores and invertivores were caught in traps at 0-100 m depth in all management areas (with some few carnivores caught even deeper, with the highest CPUE, kg/hrs, shallower than 50 m depth (Figure 7). Herbivores had a more restricted range, with no catches made deeper than 65 m bottom depth. There were only two catches of corallivores, the threadfin butterflyfish *Chaetodon auriga*, one in area 2 at 55 m depth and the other in area 6 at 17 m bottom depth. Planktivores were generally caught near the surface, shallower than 25 m, except for one catch of *Paracaesio sordius* (Lutjanidae) in area 7 at 66 m bottom depth(Figure 7).

**Figure 7.**
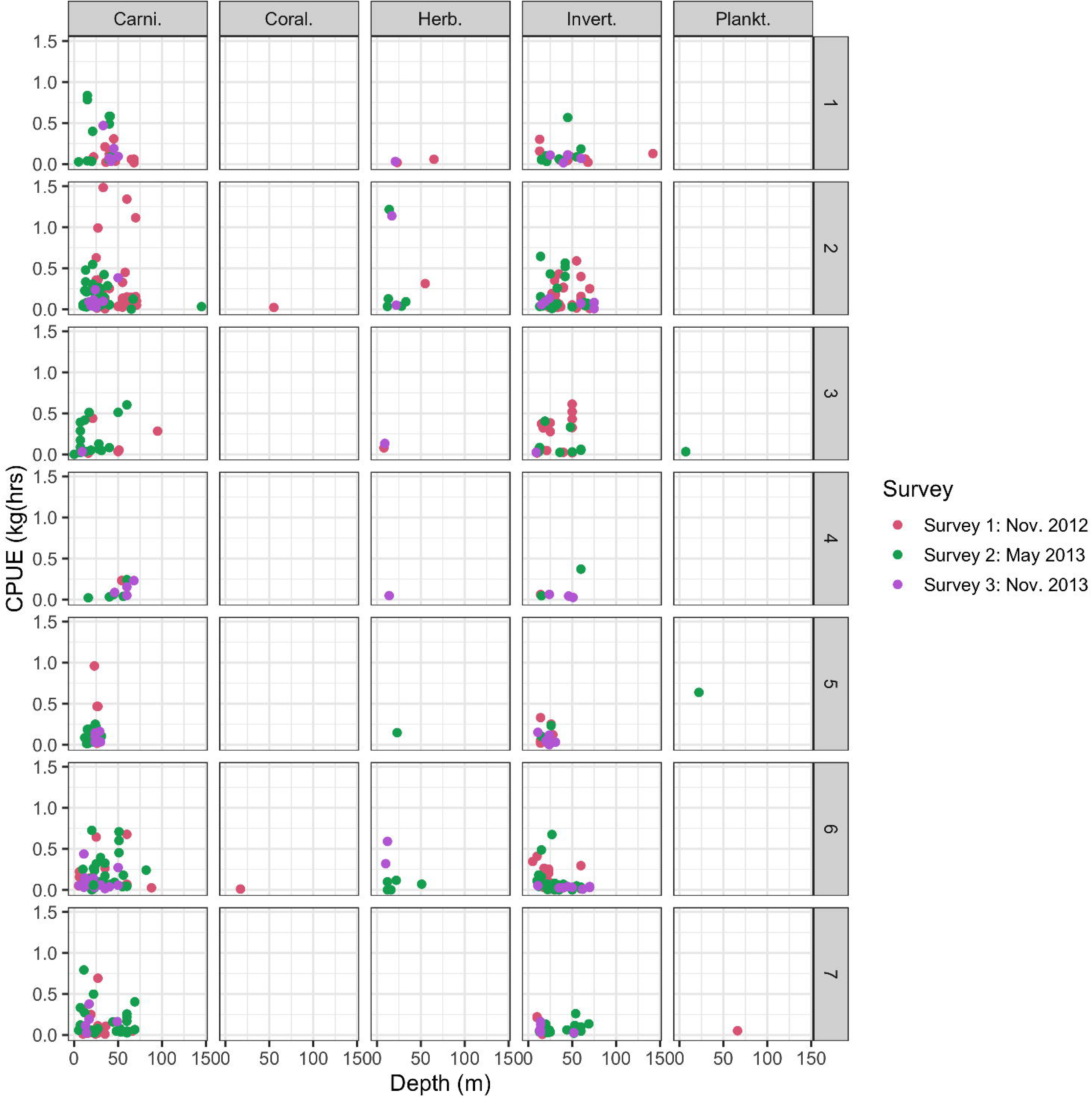
Catch per unit effort (kg/hrs fishing) of carnivores (carni), corallivores (Coral.), herbivores (Herb.), invertivores (Invert.) and planktivorous (Plankt.) fish, versus depth for traps set during the three surveys (Nov. 2012, May 2013 and Nov. 2013) in each of the seven management areas (see. Figure 1).

#### 3.2.4 Trophic level of trap catches

The mean trophic level in the catches was 3.84, with averages per area and survey ranging from 3.20 (area 3, Nov. 2013 survey) to 3.97 (area 3, May 2013 survey) (Figure 8), although when averaged across surveys the range in trophic level narrowed to 3.76 in area 6 to 3.83 in area 1 (Table 2). Trophic level per area and survey was dependent on balance in occurrence between trophic groups as the carnivores have a higher trophic level than the invertivores, the two most common trophic groups in the trap catches.

**Figure 8.**
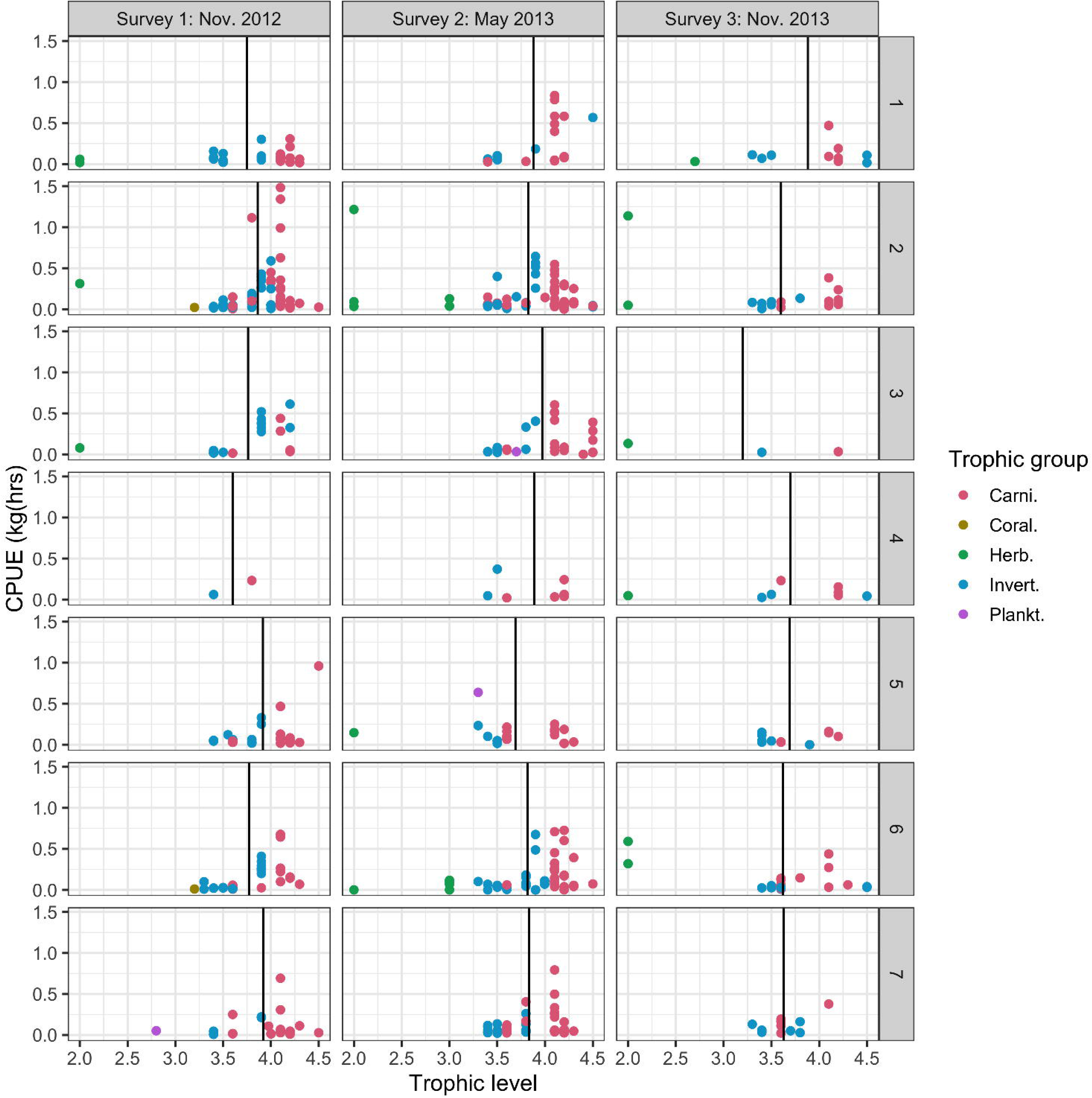
Catch per unit effort (kg per hour of fishing) versus trophic level for traps by each management area and survey, with the trophic groups: carnivores (Carni.), corallivores (Coral.), herbivores (Herb.), invertivores (Invert.) and planktivores (Plankt.) given different coloured symbols. Black vertical line shows the mean trophic level of the trap catches in each respective area and survey panel-plot.

#### 3.2.5 Other functional traits of the trap catches: placement in the water column, gregariousness and diel activity

The majority of the trap catches were of species classified as demersal, which were caught in all areas and during all surveys. These were dominating the catches except in area 3 during survey 2 where benthic species were most common (Figure 9). During the November 2013 survey there was no station with catches other than demersal fish, while both benthic and pelagic species were caught in the other two surveys, benthic fish being the second most common group after the demersals. The most dominant family groups of demersals were the Lutjanidae (mainly *Lutjanus bohar* and *L. gibbus)* and Letrhinidae (mainly *Lethrinus lentjan* and *L. mahsena*), while the benthic group was dominated by the moray eel *Gymnothorax javanicus*. Carangidae (most common species being *Carangoides bajad*) and Scombridae (most common species *Scomberoides lysan*) were the two most common families in the “Non-site attached pelagic” group.

**Figure 9.**
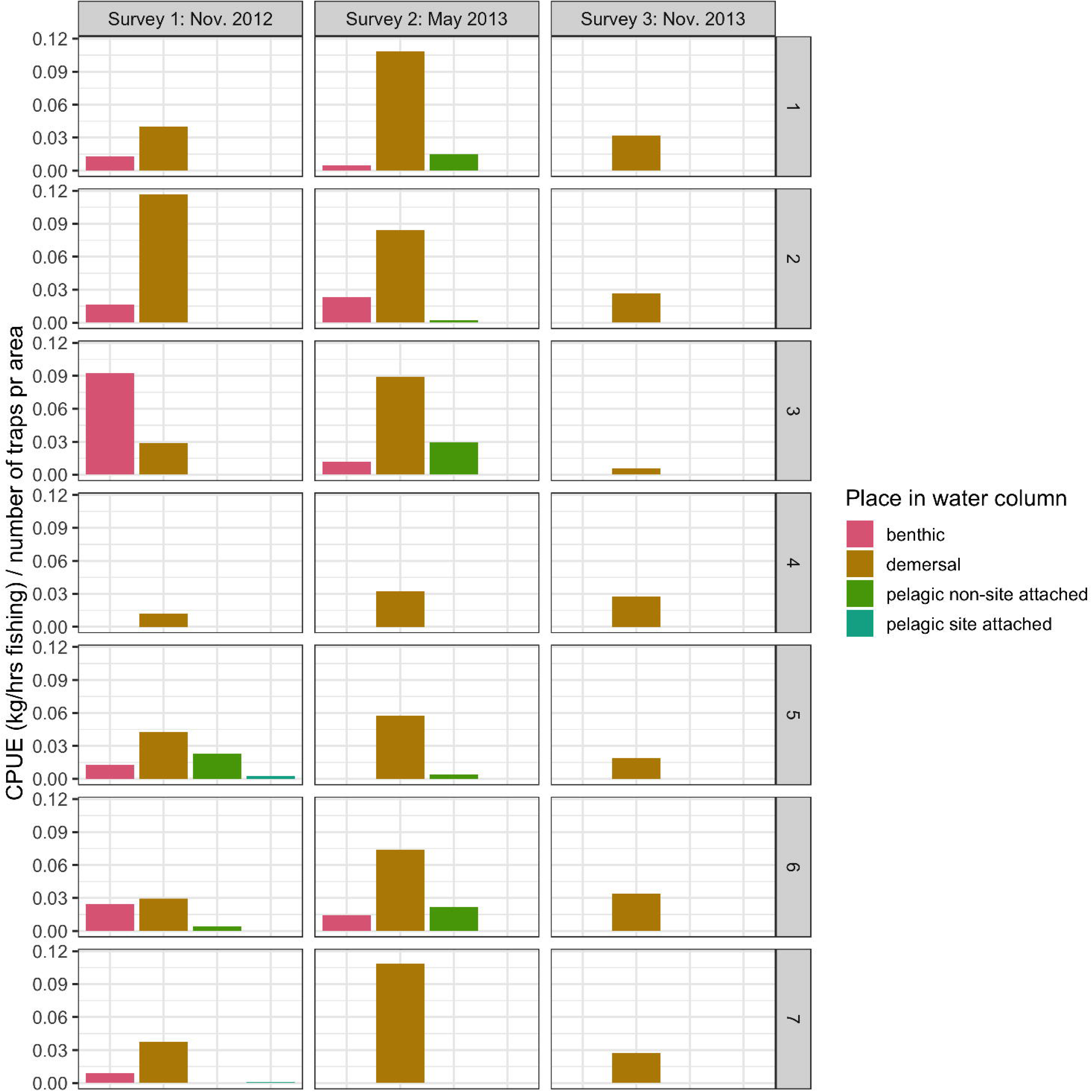
Cumulative catch rates (kg per hour of fishing) per trap for each management area and survey by the species place in the water column: benthic, demersal, pelagic non-site attached, and pelagic site attached

Day-active fish dominated the catches in all areas and surveys. Night active fish were hardly caught during the November 2013 survey, but were prevalent in the November 2012 survey particularly in area 3 (SI Figure 1).

Solitary fish (gregarious level 1) were the most common in the trap catches, except in areas 6 and 7 during the May and November 2013 surveys and in area 4 during the November 2012 survey where species of gregarious level 2 were most abundant. Species of gregarious level 3 were caught in the November 2012 and May 2013, and only in areas 2, 3, 5 and 7, but not at all during the November 2013 survey (SI Figure 2).

### 3.3 Species density

The highest number of species was observed in area 2 (36), followed by area 3, 6 and 7 (25 in each). Species density (number of species per hour of fishing - soak time of traps) ranged from 0.008 (areas 1, 2 and 6) to 0.019 (area 4) (Table 2). Area 4 had the highest species density across all surveys, while the area with the lowest species density varied by survey: area 1 in November 2012, area 7 in May 2013, and area 2 in November 2013 (Figure 10). The variability and relative difference in rank of species diversity per area was confirmed by the zero-adjusted GAM model (AIC: - 4161, df: 12), where all surveys were significant factors in the model. In addition, area 1, 2, 3 and 4 were also significant factors affecting species density, while the non-significant factors were depth, and areas 5, 6 and 7 (see SI Table 2 for details).

**Figure 10.**
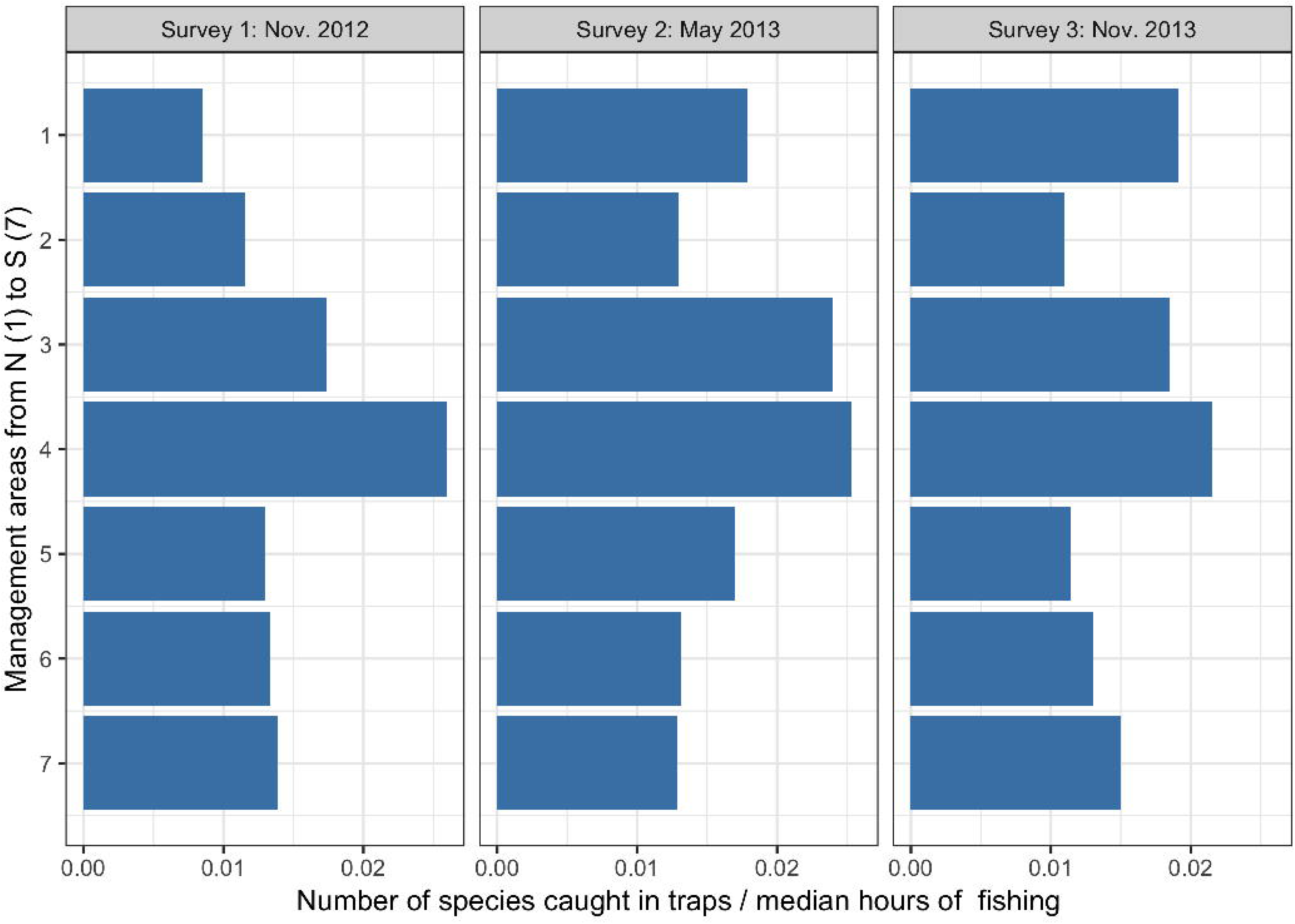
Average species densities for each of the seven management areas (number of species in trap per hour of fishing), by the three surveys.

### 3.4 Functional diversity

The Rao’s Q functional group richness index estimated using the functional diversity model excluding the ‘max length’ trait was highest (0.102) in area 2 and lowest (0.050) in area 6 (Table 2). The second-highest Rao’s Q index was found in area 5 (0.085). This was similar to the Rao’s Q index calculated from the model including all traits were area 2 had the highest and area 6 the lowest Rao’s Q index.

## 4 Discussion

The three surveys conducted in 2012 and 2013 constitute the first baseline monitoring of harvested fish along the Sudanese Red Sea coast. The first survey in November 2012 was essentially a pilot survey testing the efficiencies, and performance of various fishing gear in this region. There were significant statistical differences in both catch rates, species diversity and functional diversity between the three surveys, with the November 2013 survey being most limited, both in terms of sampling intensity and geographic coverage in the southern region (Table 1) and sampling density in area 3 compared to the previous two surveys (Figure 2).

Our sampling aimed at covering representative reefs along the entire coast at varied depths. As the species composition of coral reefs is known to be depth-stratified we intended to use gillnets to sample the surface layer, and traps and handlines to sample from the surface to 200 m. However, sampling was effectively limited by bottom conditions, e.g., ledges, and avoiding entanglement in corals. This proved challenging as the seaward side of the reefs typically had steep drop-offs to depths, often exceeding 200 m, while the areas between reefs or towards shore typically had depths of less than 100 m. Consequently, few traps were set at depths deeper than 75 m (Figure 3). Nevertheless, we achieved a reasonably comparable coverage of depths in all management areas, except for in area 5 (all surveys) and area 4 (Nov. 2012 survey) (Figure 3), thus avoiding biasing our analysis of catch rates and species diversity by differences in depth. This is further confirmed by the GAM models of CPUE and species density (S1 Table 2) where depth was not found to have a significant effect on neither CPUE nor species density.

The present analysis took a practical fisheries approach on catch per unit effort, species densities and functional diversity, focussing on the differences between the seven fisheries management areas rather than biogeographic approaches evaluating effects of distance to shore used in studies of Saudi Arabian reefs [7, 20].

Our results confirmed the presence of species already known to inhabit the Sudanese coast, although the methods used inevitably selected catchable fish above certain sizes and particular species due to the selective properties of the fishing gear, i.e. mesh selectivity (escapement, gilling, entanglement), hooks (minimum and maximum target sizes) and bait (species- and size dependent preferences and behaviour). With trap stations constituting 89% of our stations across all three surveys, it was unsurprising that the major trophic group caught were carnivores (Figure 6) as these were the species most likely to be attracted to bait. Our trap-based method was thus sub-optimal to survey herbivorous fish species such as parrotfish and most surgeonfish, and fish closely associated with coral reef habitats were most likely considerably underestimated in the catches. For such species, underwater visual census (UVC) methods, or baited remote under-water video (BRUVs) remain the only current alternatives, albeit outside the scope of this study. However, for certain vagile species such as snappers (Lutjanidae), emperors (Lethrinidae) and Scombridae the use of baited traps proved appropriate, filling a gap where handlines and gillnets were proven inefficient (see Figure 4).

The differences in catch composition compared to the artisanal fishery that targets reef-dwelling species like groupers was apparent [23], with snappers (Lutjanidae) dominating the trap catches (Figure 4). This illustrates a potential for development of fisheries targeting the snappers, emperors and other species roaming between reefs by changing the current artisanal fishery to other gear types like traps and fishing locations. Similarly, our gillnet catches caught a markedly different species composition than handline and traps, that were more similar (Figure 4). For Serranidae, Scombridae, Lutjanidae and Lethrinidae, the differences between gillnet, traps and handlines were particularly apparent. Serranidae, the family containing the most sought-after target species in the artisanal fishery, was only somewhat prevalent in handline catches from the May 2013 survey (Figure 4). This coincides with the spawning season of the commercially important *Plectropomus* spp. [18, 34], and related species. That the presence of Serranidae in handline catches is almost exclusive to this survey is likely due to increased vulnerability to harvesting in the time of spawning and -aggregation. Some species of Scombridae were only caught by gillnet which was also the most efficient gear for capturing carangids. Some Lutjanidae (e.g. *Lutjanus bohar*) were, however, caught on both gillnets, traps as well as handlines. This is to be expected given that gillnets mostly fished the upper part of the water column, while traps and handlines fished at or near the bottom, the two gears thus targeting different habitats. Similarly, from a trophic perspective this makes sense as planktivorous fish typically are pelagic, as is seen from our trap catches where planktivores were only caught in traps set shallower than 65m bottom depth (Figure 7), although it was initially surprising that planktivores were indeed caught in baited traps. The multi-gear approach was indeed chosen precisely for this reason – to sample and investigate different depth habitats of the same reef ecosystem.

### 4.1 Catch-per-unit-effort

CPUE varied significantly by gear, area and survey, but not by depth (Table 2, and S1 Table 1). The depth ranges of our stations fished were limited, with most traps set shallower than 50 m bottom depth, which probably explains why depth was not a significant factor in the model. Gillnet CPUE was higher than that of traps in all areas, and higher in the northernmost areas (1 and 2) compared to the southernmost areas (6 and 7), also with a higher standard deviation (0.482 vs 0.033). This indicated a possible higher abundance of pelagic fish along the northern part of the coast, compared to the south.

Our findings of the lowest trap CPUE in the Port Sudan and Suakin areas (4 and 5) are similar to Klaus et al. [12] who also found the lowest fish abundance in the Suakin area. The Dungonab area (2) had the highest CPUE for traps in addition to the highest diversity of trophic groups in the catches (Figure 6) and the highest functional diversity (Rao’s Q index - see Table 2), indicating that this is the most productive region north of Port Sudan. It is also the area to the north with the widest shelf and largest shallow-water region, also bordering on the very shallow Dungonab Bay with relatively higher salinity [24]. The Dungonab Bay is a designated marine protected area covering most of management area 2, granting access to local fishers only, probably reducing fishing pressure in this area. However, several previously known spawning aggregations for *Plectropomus* spp. in this area are considered lost due to fishing [34]. Area 3, just south of Dungonab bay, had the second-highest trap CPUE (0.136) and functional diversity (Rao’s Q = 0.074) pointing to a possible ecological linkage of fish communities between these two areas.

The two southernmost areas had higher trap CPUE than the Suakin and Port Sudan areas (4 and 5), although lower than in the northernmost areas, corroborating Kattan et al. [8] who found a positive relationship between top predator biomass and distance to the nearest port. Kattan et al. [8] hypothesized that fishing pressure diminishes with distance to port as fishermen prefer close and more near-shore reefs over more distant off-shore reef areas. However, these results are not consistent, exemplified by the high CPUE estimated for Dungonab (area 2). This may, however, be explained by the higher level of protection and management in this area compared to other inshore areas. Nevertheless, our results, taken together with the results of Klaus et al. [12] and Kattan et al. [8], do indicate that the local artisanal fisheries have impacts on local fish stocks.

In contrast to Klaus et al. [12], who noted the absence of large snappers, groupers and emperors in the UVCs conducted, the present surveys found these species in abundance (although large individuals were relatively rare). This can be explained by the more extensive geographic coverage, the greater depth of the sampling gear compared to UVC depths, and larger number of sampling stations in our study than in the Klaus et al. [12] study. It can also be explained by the difference in catchability of fishing gear compared to the UVC survey employed by Klaus et al. [12]. Snappers and emperors, and to a lesser degree groupers, showed a roving behaviour between reefs when observed during dives and snorkelling, often keeping a distance from divers (personal observations), similar to what Colton and Swearer [35] observed for mobile predators, possibly making them less available in UVC transect paths. These species were, however, attracted to baited traps, explaining their common occurrence in catches in the present study.

### 4.2 Species density and functional diversity

There were differences in species densities between the seven management areas (Table 2). The highest densities was found in area 4 (Port Sudan), which also was significantly different from all other areas. The species density results are thus in contrast with Kattan et al.’s [8] findings of relatively higher biodiversity in remote areas of the coast. Our present analysis was limited by the low number of stations in several areas and surveys, in particular area 4. When evaluating the functional diversity, a different picture emerges. Area 2 had the highest Rao’s Q value of all the areas (0.102), while area 4 had a lower Rao’s Q of 0.063 in the model excluding the maximum length trait. If maximum length was included in the model area 4 had the lowest Rao’s Q value of all areas. Functional diversity (Rao’s Q) was calculated using CPUE as the abundance data, thereby weighing the species occurrence not just on hours fishing, but also by the catch weight, contrasting species densities where the species occurrence were weighted only by the fishing time. Since the CPUE in area 2 on average was higher than in area 4, except during the November 2013 survey (Figure 5), this is the most likely explanation of the discrepancies between the species densities and functional diversity. Stuart-Smith et al [30] argues that since the ecological effects of a species generally are proportional to its abundance, abundance-weighted functional diversity (e.g. Rao’s Q) more accurately reflects the functional structure of a community than diversity metrics based on simple counts or species inventories. We therefore placed higher emphasis on the Rao’s Q indices in the present analysis than on species density. The higher Rao’s Q in the Dunognab area (area 2) and the low Rao’s Q in area 4 indicate that the protected Dungonab bay area has a higher functional biodiversity than the Port Sudan area (area 4) which is the most heavily impacted by human activities along the Sudanese coast.

The lack of clear gradients in species distribution or geographically unique species indicates that the species caught in our sampling gear have fairly uniform distributions along the Sudanese coast. This could be expected given the lack of species gradients observed elsewhere in the Red Sea [3, 7]. It is inevitable that the capture-based methodologies employed to cover the entire coast during surveys has resulted in missing locally rare or endemic species. There are other issues pertaining to gillnet and trap fishing in coral reef areas that make them less desirable from a biodiversity and fisheries conservation science perspective, such as ghost nets/traps, and bycatch of illegal and/ or vulnerable species (e.g., sharks). Selective passive gears, like traps can, however, be employed with less environmental impact or bycatch of threatened elasmobranch species than pelagic gillnets or long-lines, while traps without bio-degradable openings may cause ghost fishing if lost.

Klaus et al. [12] identified the 70 km coastal region between Port Sudan and Suakin as being the most heavily affected by coastal and harbour developments and contended the notion that this had affected the reefs in this area. This is further corroborated by Kattan et al. [8], who found that biomass and species richness decreased with distance from the main port of Port Sudan. However, distance to Port Sudan is a misleading measure of the distance fishers have to travel to catch fish as they operate out of numerous landing sites all along the coast. A better measure of distance to fishing areas would be to measure the distance of the actual fishing trips from the local landing sites. However, our results did show low species density and functional diversity (Rao’s Q), and a lack of herbivores and planktivores in the Port Sudan management area and in the Arakia area just to the north, supporting the hypothesis that increased urban development and proximity to population centres have resulted in a mining out of catchable fish biomass and reduced the productivity of reefs. In management area 3 and 4 the higher human populations and number of fishermen based in these regions have likely increased fishing pressures more than in other areas, thus representing two highly likely factors in explaining the low species diversity and lower catch rates in these regions.

### 4.3 Recommendations

Whether traps are more appropriate than visual census methods, which may underestimate species that actively avoid divers doing the census [35, 36], or are reluctant to approach a baited camera rig during the relatively short recording time, remains to be properly tested for species typically targeted by Sudan’s artisanal coral reef fishery. In a study comparing the relative efficiency of commercial fish traps and BRUVs in sampling tropical demersal fishes in Western Australia, Harvey et al. [37] found that BRUVs had greater statistical power to detect changes in abundance than an equivalent number of traps. Among five commercially important Indo-Pacific species (*Epinephelus bilobatus*, *Epinephelus multinotatus*, *Lethrinus punctulatus*, *Lutjanus russelli* and *Lutjanus sebae*) only emperor red snapper (*L. sebae*) was more efficiently sampled with commercial traps. However, a monitoring system based on traps requires lower skill levels and less infrastructure than UVC- and BRUV methodology, and most species will survive capture and subsequent release if a non-extractive approach to monitoring is desired. Traps also have their drawbacks in being bulky, requiring a winch to haul and will involve more sea time if soaked overnight. Evaluating such practical constraints is essential when planning and designing fish monitoring programs in least developed countries with poor institutional capacities and limited resources like Sudan. Still, our results show the merit of a multi-methods approach to monitoring in a complex coral reef ecosystem area like the Red Sea.

### 4.4 Conclusions

We here present the first baseline survey of the fisheries resources along the Sudanese Red Sea coast, providing novel insights into the distributions, catch rates, species densities and functional diversities of fish communities along the Sudanese coast, supplementing the rather scant biogeographic information available on fish species in this area Our results demonstrate differences in CPUE, highlighting the Dungonab Bay area as a hot spot. The methods used do, however, not provide a full census of all fish species in the areas and the surveys did not cover all habitats. The methods presented should therefore be further developed and complemented with visual census-based methods to cover the full diversity, and include all functional groups. Such a complementary approach may yield further improved assessment of the fish distribution, abundance and species richness in the highly complex coral-reef environment of the Sudanese coast.

The observed species densities, functional diversity and catch rates demonstrate clear local (Figure 8) and seasonal variabilities (Figure 6), as well differences in CPUE between management regions that can be hypothesised to be caused by varying degrees of human impacts along the coast of Sudan. With an increasing human population, increasing coastal development and a push to expand and increase fisheries, sustainable and ecosystem-based management plans should be developed, implemented and enforced to avoid overfishing, habitat destruction and the associated negative socioeconomic impact on the poor and fragile fishing communities along the coasts of the Red Sea. The present study provides a first baseline to use as foundation for developing such plans.

## Acknowledgements

We wish to acknowledge the support from the Sudanese partner institutions: The Marine Fisheries Administration, The Fisheries Research Institute and the University of the Red Sea State – Faculty of Marine Science. Asbjørn Aasen of the IMR is thanked for designing and building the collapsible traps used in these surveys. The scientific and technical crew on the ‘Don Questo’ are thanked for their dedicated work with sample collection. Rick Stuart Smith is thanked for providing a table of biological traits for coral reef fish species. Funding was provided by the Norwegian Government through the Embassy in Khartoum, and the project was administrated by UNIDO.

## References

1. Berumen ML, Hoey AS, Bass WH, Bouwmeester J, Catania D, Cochran JEM, et al. The status of coral reef ecology research in the Red Sea. Coral Reefs. 2013;32: 737–748. doi:10.1007/s00338-013-1055-8

2. DiBattista JD, Howard Choat J, Gaither MR, Hobbs JPA, Lozano Cortés DF, Myers RF, et al. On the origin of endemic species in the Red Sea. Journal of Biogeography. 2016;43: 13–30. doi:10.1111/jbi.12631

3. DiBattista JD, Roberts MB, Bouwmeester J, Bowen BW, Coker DJ, Lozano Cortés DF, et al. A review of contemporary patterns of endemism for shallow water reef fauna in the Red Sea. Journal of Biogeography. 2nd ed. 2016;43: 423–439. doi:10.1111/jbi.12649

4. Kessel ST, Elamin NA, Yurkowski DJ, Chekchak T, Walter RP, Klaus R, et al. (2017) Conservation of reef manta rays (Manta alfredi) in a UNESCO World Heritage Site: Large-scale island development or sustainable tourism? PLoS ONE 12(10): e0185419. https://doi.org/10.1371/journal.pone.0185419

5. Loya Y, Genin A, el-Zibdeh M, Naumann MS, Wild C. Reviewing the status of coral reef ecology of the Red Sea: key topics and relevant research. Coral Reefs. 2014;33: 1179–1180. doi:10.1007/s00338-014-1170-1

6. Tesfamichael D, Pauly D. Introduction to the Red Sea. The Red Sea Ecosystem and Fisheries. Dordrecht: Springer, Dordrecht; 2016. pp. 1–19. doi:10.1007/978-94-017-7435-2_1

7. Roberts MB, Jones GP, McCormick MI, Munday PL, Neale S, Thorrold S, et al. Homogeneity of coral reef communities across 8 degrees of latitude in the Saudi Arabian Red Sea. Marine Pollution Bulletin. 2016;105: 558–565. doi:10.1016/j.marpolbul.2015.11.024

8. Kattan A, Coker DJ, Berumen ML. Reef fish communities in the central Red Sea show evidence of asymmetrical fishing pressure. Mar Biodiv. Springer Berlin Heidelberg; 2017;6: 1–12. doi:10.1007/s12526-017-0665-8

9. DuTemple LA. Jacques Cousteau. Minneapolis: Lerner Publications Company; 2000.

10. Randall JE. The divers guide to Red Sea reef fishes. London; 1996.

11. Debelius H. Red Sea reef guide. 5 ed. Frankfurt: IKAN; 2011.

12. Klaus R, Kemp J, Samoilys M, Anlauf H, Din El S, Abdalla EO, et al. Ecological patterns and status of the reefs of Sudan. 2009. pp. 1–5.

13. Bundy A, Chuenpagdee R, Boldt JL, de Fatima Borges M, Camara ML, Coll M, et al. Strong fisheries management and governance positively impact ecosystem status. Fish and Fisheries. 2016;18: 412–439. doi:10.1111/faf.12184

14. Pauly D, Zeller D. Accurate catches and the sustainability of coral reef fisheries. Current Opinion in Environmental Sustainability. Elsevier B.V; 2014;7: 44–51. doi:10.1016/j.cosust.2013.11.027

15. Newton K, Côté IM, Pilling GM, Jennings S, Dulvy NK. Current and future sustainability of island coral reef fisheries. Current Biology. 2007;17: 655–658. doi:10.1016/j.cub.2007.02.054

16. Johnson AE, Cinner JE, Hardt MJ, Jacquet J, McClanahan TR, Sanchirico JN. Trends, current understanding and future research priorities for artisanal coral reef fisheries research. Fish and Fisheries. Blackwell Publishing Ltd; 2013;14: 281–292. doi:10.1111/j.1467-2979.2012.00468.x

17. Nash KL, Graham NAJ. Ecological indicators for coral reef fisheries management. Fish and Fisheries. 2016;17: 1029–1054. doi:10.1111/faf.12157

18. Elamin S. Stock assessment and population dynamics of Plectropomus pessuliferus and Plectropomus areolatus in the Sudanese red sea coast. Terengganu University. 2012.

19. Spaet JLY, Berumen ML. Fish market surveys indicate unsustainable elasmobranch fisheries in the Saudi Arabian Red Sea. Fisheries Research. 2015;161: 356–364. doi:10.1016/j.fishres.2014.08.022

20. Khalil MT, Bouwmeester J, Berumen ML. Spatial variation in coral reef fish and benthic communities in the central Saudi Arabian Red Sea. PeerJ. PeerJ Inc; 2017;5: e3410. doi:10.7717/peerj.3410

21. Trewavas E. Nomeclature of the Tilapias of southern Africa. Journal of the Limnological Society of Southern Africa. Food and Agriculture Organisation of the United … 2010;7: 42–42. doi:10.1080/03779688.1981.9632937

22. FAO. The Republic of Sudan. In: fao.org [Internet]. [cited 1 Oct 2017]. Available: http://www.fao.org/fishery/facp/SDN/en

23. Tesfamichael D, Elawad AN. Sudan. The Red Sea Ecosystem and Fisheries. Dordrecht: Springer, Dordrecht; 2016. pp. 37–48. doi:10.1007/978-94-017-7435-2_3

24. Olsen O, Wenneck W, Aasen A, Fadula F, Abdalla Nasir A, Habiballah H. First survey of renewable marine resources in the Red Sea State, Republic of Sudan 1 – 30 Nov 2012. 2015.

25. Froese R, Pauly D. FishBase 2000 Concepts, design and data sources. Froese R, Pauly D, editors. Los Banos, Laguna, Phillipines: ICLARM; 2000 p. 344.

26. Gabr MH, Mal AO. Trammel net fishing in Jeddah: species composition, relative importance, length-weight and length-girth relationships of major species. International Journal of Fisheries and Aquatic Studies. 2018;6: 305–313.

27. Lewis M, Jordan S, Chancy C, Harwell L, Goodman L, Quarles R. Summer Fish Community of the Coastal Northern Gulf of Mexico: Characterization of a Large-Scale Trawl Survey. Transactions of the American Fisheries Society. 2007;136: 829–845. doi:10.1577/T06-077.1

28. Azarovitz TR. A brief historical review of the Woods Hole Laboratory trawl survey time series. Canadian Special Publication of Fisheries and Aquatic Sciences. 1981;58: 62–67.

29. Harley SJ, Myers RA, Dunn A. Is catch-per-unit-effort proportional to abundance? Can J Fish Aquat Sci. NRC Research Press Ottawa, Canada; 2011;58: 1760–1772. doi:10.1139/f01-112

30. Stuart-Smith RD, Bates AE, Lefcheck JS, Duffy JE, Baker SC, Thomson RJ, et al. Integrating abundance and functional traits reveals new global hotspots of fish diversity. Nature. Nature Publishing Group; 2013;501: 539–542. doi:10.1038/nature12529

31. Stroemme T. NAN-SIS: software for fishery survey data logging and analysis: user’s manual. Computerized Information Series: Fisheries (FAO). FAO; 1992.

32. Team RC. R Language Definition. 2000;: 1–60.

33. UNESCO. Nomination of Sanganeb Marine National Park And Dungonab Bay/Mukkawar Island Marine National Park (Sudan – Red Sea). whc.unesco.org. Khartoum; 2016 Jan p. 461.

34. Elamin, S.M. 2017. Final report on the first assessment and monitoring programme for Plectropomus spp. “Najil” in Dungonab National Marine Park. Faculty of Marine Science and Fisheries, Red Sea University, Port Sudan.

35. Colton MA, Swearer SE. A comparison of two survey methods: differences between underwater visual census and baited remote underwater video. Mar Ecol Prog Ser. 2010;400: 19–36. doi:10.3354/meps08377

36. Emslie, M.J., Cheal, A.J., MacNeil, M.A., Miller, I.R., Sweatman, H.P.A. 2018. Reef fish communities are spooked by scuba surveys and may take hours to recover. PeerJ 6:e4886; doi:10.7717/peerj.4886

37. Harvey, E.S., Newman, S.J., McLean, D.L., Cappo, M., Meeuwig, J.J., Skepper, C.L. 2012. Comparison of the relative efficiencies of stereo-BRUVs and traps for sampling tropical continental shelf demersal fishes. Fisheries Research 125–126: 108-120 doi: 10.1016/j.fishres.2012.01.026

