## Supplementary materials for "Distribution and diversity of fish species exposed to artisanal fishery along the Sudanese Red Sea coast"

Table 1 Output of zero-inflated GAM model analysis of biomass CPUE of traps (TB), handlines (HL) and gillnets (intercept) as a function of depth, management area, survey and fishing gear. Analysis done in R version 3.5.2 (Eggshell Igloo) using the ‘gamlss’ package (see [www.gamlss.com](http://www.gamlss.com) for details)

```
## ******************************************************************
## Family:  c("ZAGA", "Zero Adjusted GA")
##
## Call:  gamlss(formula = CPUEw ~ depth + factor(id) + factor(survey) +
##     factor(gear), family = ZAGA, data = c2)
##
## Fitting method: RS()
##
## ------------------------------------------------------------------
## Mu link function:  log
## Mu Coefficients:
##                        Estimate Std. Error t value Pr(>|t|)
## (Intercept)           -1.183870   0.181345  -6.528 9.87e-11 ***
## depth                  0.003163   0.002327   1.359 0.174344
## factor(id)2           -0.066567   0.149632  -0.445 0.656495
## factor(id)3           -0.076983   0.202036  -0.381 0.703246
## factor(id)4           -0.068939   0.202976  -0.340 0.734189
## factor(id)5           -0.513933   0.167731  -3.064 0.002233 **
## factor(id)6           -0.401688   0.162065  -2.479 0.013331 *
## factor(id)7           -0.585171   0.164508  -3.557 0.000390 ***
## factor(survey)2013002 -0.162780   0.095859  -1.698 0.089748 .
## factor(survey)2013005 -0.224448   0.111011  -2.022 0.043418 *
## factor(gear)HL         0.775526   0.194477   3.988 7.08e-05 ***
## factor(gear)TB        -0.415457   0.108136  -3.842 0.000129 ***
## ---
## Signif. codes:  0 '***' 0.001 '**' 0.01 '*' 0.05 '.' 0.1 ' ' 1
##
## ------------------------------------------------------------------
## Sigma link function:  log
## Sigma Coefficients:
##             Estimate Std. Error t value Pr(>|t|)
## (Intercept)  0.08571    0.02157   3.974 7.49e-05 ***
## ---
## Signif. codes:  0 '***' 0.001 '**' 0.01 '*' 0.05 '.' 0.1 ' ' 1
##
## ------------------------------------------------------------------
## Nu link function:  logit
## Nu Coefficients:
##             Estimate Std. Error t value Pr(>|t|)
## (Intercept) -0.74653    0.06204  -12.03   <2e-16 ***
## ---
## Signif. codes:  0 '***' 0.001 '**' 0.01 '*' 0.05 '.' 0.1 ' ' 1
##
## ------------------------------------------------------------------
## No. of observations in the fit:  1191
## Degrees of Freedom for the fit:  14
##       Residual Deg. of Freedom:  1177
##                       at cycle:  2
##
## Global Deviance:     279.0767
##             AIC:     307.0767
##             SBC:     378.2324
## ******************************************************************
```

Table 2 Output of zero-inflated GAM model analysis of species density (number of species pr hour of fishing) for traps as a function of depth, management area and survey. Analysis done in R version 3.5.2 (Eggshell Igloo) using the ‘gamlss’ package (see [www.gamlss.com](http://www.gamlss.com) for details)

```
## ******************************************************************
## Family:  c("ZAGA", "Zero Adjusted GA")
##
## Call:  gamlss(formula = Species_Hrs ~ depth + factor(area) +
##     factor(survey), family = ZAGA, data = spue.df)
##
## Fitting method: RS()
##
## ------------------------------------------------------------------
## Mu link function:  log
## Mu Coefficients:
##                         Estimate Std. Error  t value Pr(>|t|)
## (Intercept)           -3.0456597  0.0282803 -107.695  < 2e-16 ***
## depth                  0.0003143  0.0003925    0.801 0.423536
## factor(area)2          0.0952550  0.0257799    3.695 0.000238 ***
## factor(area)3          0.1083585  0.0319608    3.390 0.000740 ***
## factor(area)4          0.0834147  0.0390748    2.135 0.033148 *
## factor(area)5          0.0540349  0.0296866    1.820 0.069184 .
## factor(area)6         -0.0337542  0.0278053   -1.214 0.225200
## factor(area)7         -0.0464743  0.0294473   -1.578 0.114993
## factor(survey)2013002  0.2071281  0.0174699   11.856  < 2e-16 ***
## factor(survey)2013005  0.2470968  0.0190004   13.005  < 2e-16 ***
## ---
## Signif. codes:  0 '***' 0.001 '**' 0.01 '*' 0.05 '.' 0.1 ' ' 1
##
## ------------------------------------------------------------------
## Sigma link function:  log
## Sigma Coefficients:
##             Estimate Std. Error t value Pr(>|t|)
## (Intercept) -1.66955    0.02708  -61.66   <2e-16 ***
## ---
## Signif. codes:  0 '***' 0.001 '**' 0.01 '*' 0.05 '.' 0.1 ' ' 1
##
## ------------------------------------------------------------------
## Nu link function:  logit
## Nu Coefficients:
##             Estimate Std. Error t value Pr(>|t|)
## (Intercept)    -16.2      129.2  -0.125      0.9
##
## ------------------------------------------------------------------
## No. of observations in the fit:  674
## Degrees of Freedom for the fit:  12
##       Residual Deg. of Freedom:  662
##                       at cycle:  2
##
## Global Deviance:     -4185.753
##             AIC:     -4161.753
##             SBC:     -4107.594
## ******************************************************************
```

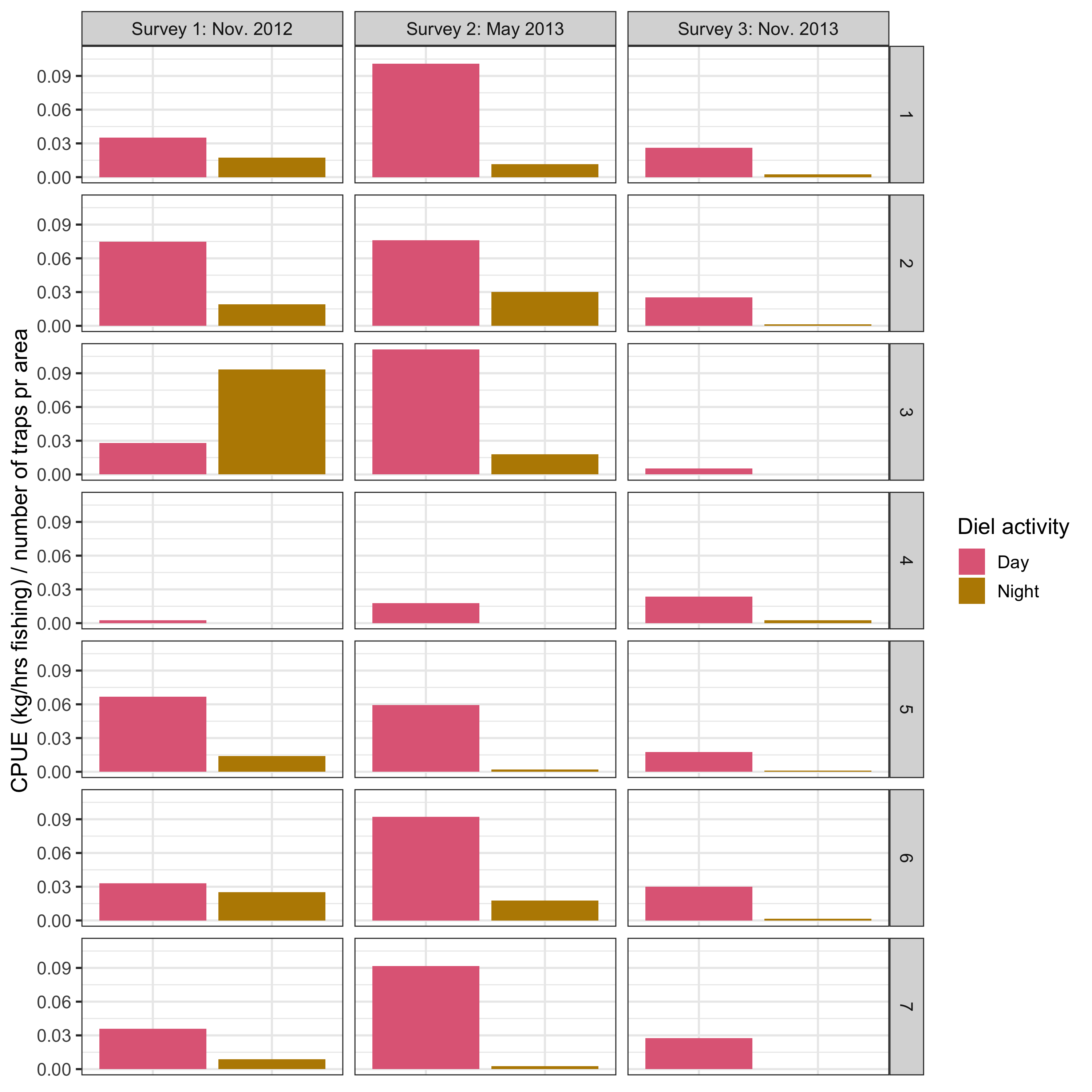

Figure 1 CPUE of traps grouped by the diel activity of the fish caught by management area and survey.

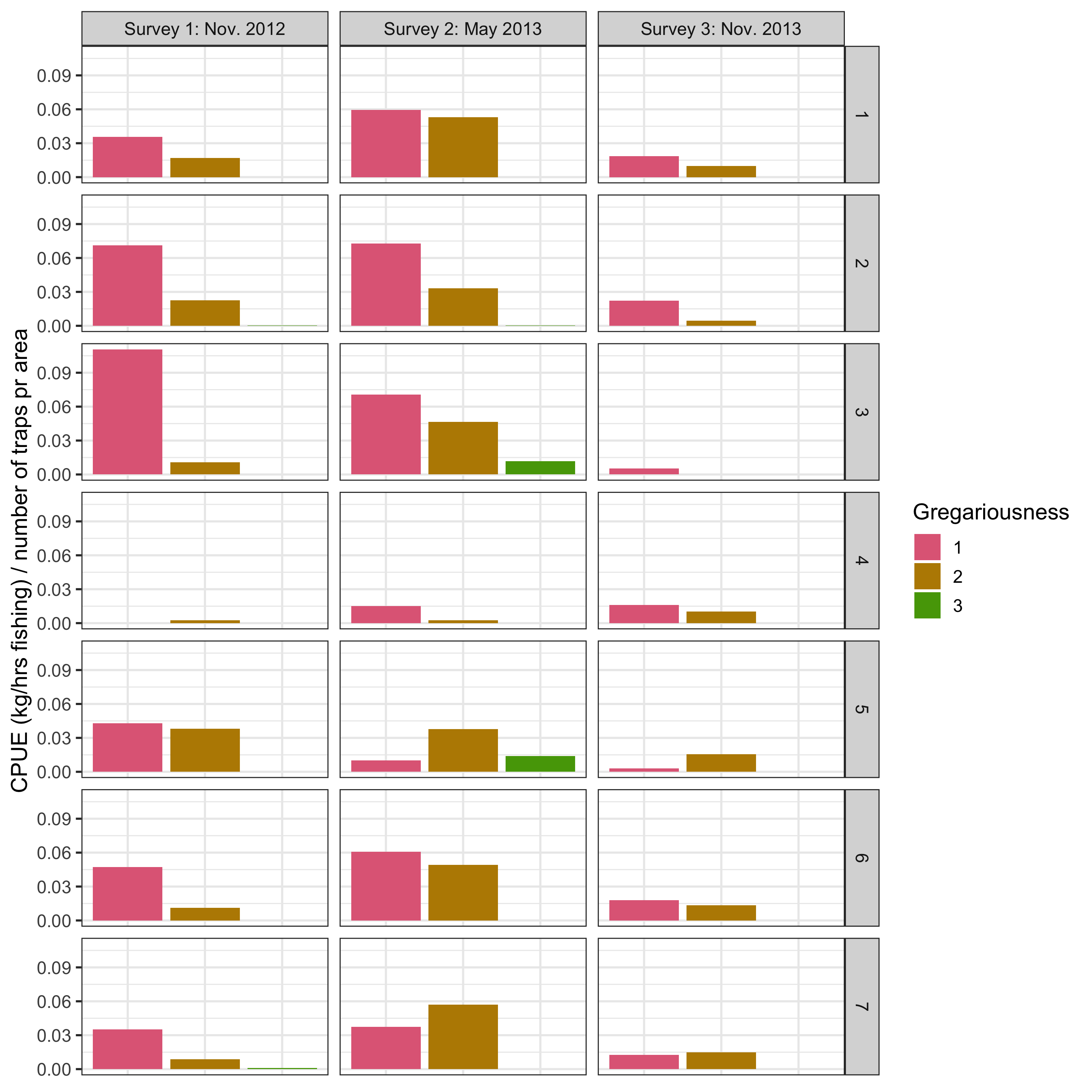

Figure 2 CPUE of traps grouped by the gregariousness (1:solitary to 3: schooling) of the fish caught by management area and survey.
